# Prot2Prop: Structure-informed multitask protein property prediction

**DOI:** 10.64898/2026.06.28.735009

**Authors:** Danial Gharaie Amirabadi, Cody Jackson, Dong Su Kim, Maximilian Sprang, Keaun Amani

**Affiliations:** Neurosnap Research, Neurosnap Inc; Department of Dermatology University Medical Center of the Johannes Gutenberg University Mainz; Institute of Quantitative and Computational Biology, Johannes Gutenberg University Mainz

## Abstract

Protein engineering often relies on separate models for related developability properties, limiting efficiency and transfer across tasks. We present Prot2Prop, a multitask framework based on a frozen ProstT5 encoder with shared and task-specific adapters for joint prediction of six protein properties: material production, solubility, temperature stability, aggregation propensity, expression yield, and folding stability. Across held-out test data, Prot2Prop achieved strong performance on both classification and regression tasks, including AUROC values ranging from 0.86 to 0.98 for classification endpoints and Spearman correlations ranging from 0.73 to 0.86 for regression endpoints. The model achieved particularly strong performance for temperature stability (AUROC = 0.98) and aggregation propensity (Spearman = 0.86). Post-hoc calibration further improved regression accuracy, reducing folding stability MAE from 0.67 to 0.48. These results demonstrate that parameter-efficient multitask adaptation of protein language models can provide accurate and unified prediction of diverse protein developability properties.

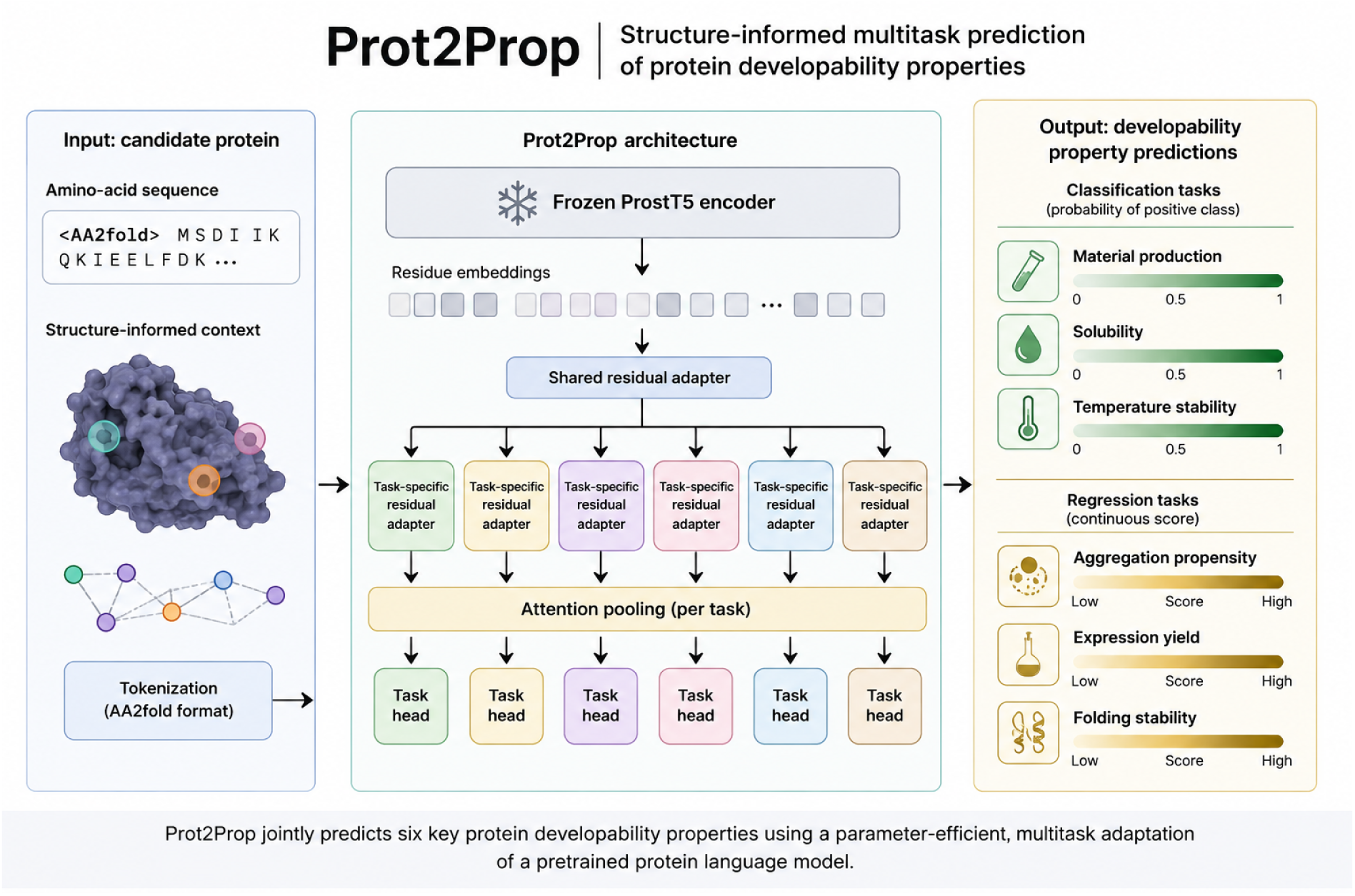

## Introduction

The ability to accurately predict protein properties such as solubility, thermal stability, aggregation propensity, expression yield, and folding stability is central to modern protein engineering, therapeutic design, and synthetic biology (Kuriata *et al*., 2019; Qing *et al*., 2022; Rosace *et al*., 2023; Pudžiuvelytė *et al*., 2024). These properties strongly influence whether a candidate protein can be expressed, purified, formulated, and deployed successfully in downstream applications (Thumuluri *et al*., 2022; Rosace *et al*., 2023; Pudžiuvelytė *et al*., 2024). In practice, however, these traits are rarely considered in an integrated manner. Instead, researchers often rely on separate task-specific predictors, each trained independently on a narrow dataset and optimized for a single endpoint (Chandra *et al*., 2023). This fragmented modeling landscape makes it difficult to evaluate proteins holistically and prevents the reuse of shared biochemical signals across related developability tasks (An and Weng, 2022).

This limitation is particularly important because many protein properties are biologically and statistically coupled (An and Weng, 2022; Qing *et al.,* 2022; Rosace *et al.,* 2023). A mutation that improves one trait, such as thermostability, may also alter folding behavior, solubility, or aggregation risk (Kuriata *et al.,* 2019; Qing *et al.,* 2022; Rosace *et al.,* 2023). Likewise, expression yield often reflects a combination of structural stability, sequence composition, and higher-order developability effects rather than a single isolated mechanism (Thumuluri *et al.,* 2022; Chen *et al.,* 2023). Treating these outcomes as independent prediction problems therefore leaves performance on the table, especially when some tasks are data-rich and others are comparatively sparse (An and Weng, 2022; Schmirler, Heinzinger and Rost, 2024). This motivation is consistent with prior work in biological prediction, where multitask neural networks have been shown to outperform single-task models by sharing information across related assays and targets at large scale (Ramsundar *et al*., 2015). More recent work in protein language model adaptation has similarly suggested that joint training across related biological objectives can improve transfer and reduce the need for separate endpoint-specific models (Anyaegbunam *et al*., 2026). A unified model that learns across multiple related endpoints has the potential to improve both parameter efficiency and predictive performance by capturing common latent determinants of protein behavior (Houlsby *et al.,* 2019; An and Weng, 2022; Schmirler, Heinzinger and Rost, 2024).

Recent protein language models have created an opportunity to revisit this problem with substantially stronger priors than were available in earlier generations of developability predictors (Brandes *et al*., 2022; Unsal *et al*., 2022; Chandra *et al*., 2023). In particular, large pretrained models such as ProstT5 provide rich sequence representations informed by protein structural regularities, enabling transfer to downstream tasks with relatively modest fine-tuning (Heinzinger *et al*., 2024; Schmirler, Heinzinger and Rost, 2024). Yet many existing applications still adapt these models in a single-task setting, requiring one model per property and offering limited flexibility when heterogeneous outputs must be predicted jointly across both classification and regression tasks (Brandes *et al*., 2022; Unsal *et al*., 2022; Schmirler, Heinzinger and Rost, 2024). This is especially restrictive in practical screening workflows, where users need one system that can score multiple properties at once rather than a collection of disconnected models with inconsistent behavior and calibration (An and Weng, 2022; Rosace *et al*., 2023).

At the same time, fully fine-tuning large protein language models for every property is computationally expensive and operationally inefficient (Houlsby *et al*., 2019; Schmirler, Heinzinger and Rost, 2024). A more scalable alternative is parameter-efficient adaptation, where the pretrained backbone remains frozen and only lightweight trainable modules are introduced for downstream specialization (Houlsby *et al*., 2019; Schmirler, Heinzinger and Rost, 2024; Sledzieski *et al*., 2024). Such an approach is appealing for multitask protein property prediction because it allows the model to retain a shared biochemical representation while learning task-specific corrections only where necessary (Houlsby *et al*., 2019; An and Weng, 2022; Schmirler, Heinzinger and Rost, 2024). However, an effective multitask system must balance shared transfer against task-level specialization, particularly when the target set spans binary classification tasks alongside continuous regression objectives with different scales, sample sizes, and label semantics (An and Weng, 2022; Schmirler, Heinzinger and Rost, 2024).

In this study, we present Prot2Prop, a parameter-efficient multitask framework for joint prediction of protein developability properties. Prot2Prop is built on a frozen ProstT5 encoder (Heinzinger *et al.,* 2024) and combines a shared residual adapter with lightweight task-specific residual adapters (Houlsby *et al.,* 2019; Schmirler, Heinzinger and Rost, 2024), attention-based sequence pooling, and task-specific prediction heads. This design enables the model to learn a common representation across tasks while allowing each endpoint to specialize its token-level features before sequence summarization and prediction. ProstT5 was selected as the backbone for two reasons. First, preliminary exploratory experiments with several candidate protein language models suggested that it provided the most promising early performance on our multitask benchmark. Because these evaluations were conducted during the initial prototyping stage and were not designed as a systematic comparative study, we do not make broader claims regarding its superiority over alternative backbones. Second, ProstT5 supports representations derived from both protein sequence and structure through its sequence-to-structure translation framework, providing flexibility for future extensions that incorporate structural information. In contrast, sequence-only language models such as ESM2 (Lin *et al*., 2023) do not natively encode structural representations in their input space. The resulting architecture supports joint modeling of heterogeneous labels within a single framework, spanning both binary and continuous outputs.

### Datasets and Evaluation Protocol

#### Data Sources

Prot2Prop was trained on six protein developability-related prediction tasks spanning both classification and regression objectives: **material production**, **solubility**, **temperature stability**, **aggregation propensity**, **expression yield**, and **folding stability**. The three classification tasks were assembled from publicly available sequence-level datasets provided through the AI4Protein (D. Wang *et al*., 2025) collection, whereas the three regression tasks were assembled from ProteinGym (Notin *et al*., 2023) deep mutational scanning datasets using task-specific manifests. In the current implementation, material production, solubility, and temperature stability are modeled as binary sequence-level classification tasks, while aggregation propensity, expression yield, and folding stability are modeled as continuous sequence-level regression tasks.

To support multitask learning, all datasets were converted into a unified sequence-centric representation in which each unique protein sequence was associated with zero or more task labels. During aggregation, each source dataset was mapped to standardized task metadata, including task name, label type, prediction head type, number of classes, and loss family. Binary classification labels were coerced to 0/1 values, whereas regression labels were retained as continuous floating-point values. After aggregation, labels for the same sequence were assembled into a multitask label vector together with a task-specific observation mask indicating which endpoints were present for that sequence. This procedure preserved the original task definitions while placing all datasets into a common format suitable for joint multitask training.

#### Data Splitting Strategy

After aggregation, unique protein sequences were partitioned at the sequence level into training, validation, and test splits using fixed global fractions of **80% / 10% / 10%.** This global split was applied after merging task labels by sequence, ensuring that the same protein sequence could not appear in multiple splits under different tasks. This design was important for multitask evaluation because many sequences were only partially annotated and contributed labels to only a subset of endpoints.

During training, proteins with incomplete task coverage were retained and losses were computed only for tasks with observed labels, allowing partially annotated sequences to contribute supervision wherever labels were available. Regression targets were normalized using statistics computed from the training split only, whereas evaluation metrics were reported in the original label space.

All model selection, architectural refinement, and seed selection for the main manuscript results were based on validation-set performance.

#### Evaluation Metrics

For classification tasks, performance was evaluated using **accuracy**, **balanced accuracy**, **precision**, **recall**, **F1 score**, **area under the receiver operating characteristic curve (AUROC)**, and **area under the precision-recall curve (AUPRC)**. Threshold-dependent metrics such as accuracy, precision, recall, and F1 were used to assess direct classification behavior, whereas AUROC and AUPRC were used to assess threshold-independent ranking quality from predicted class probabilities.

For regression tasks, performance was evaluated using **mean absolute error (MAE)**, **root mean squared error (RMSE)**, and **Spearman rank correlation**. MAE and RMSE quantify the magnitude of prediction error, whereas Spearman correlation measures preservation of rank ordering between predicted and observed values. Because several regression endpoints showed strong rank fidelity despite residual scale or offset bias, both error-based and rank-based metrics were retained throughout evaluation.

#### Post-hoc Calibration Analysis

In addition to raw model outputs, we evaluated simple post-hoc calibration procedures for both classification and regression tasks. For binary classification endpoints, decision thresholds were tuned from predicted positive-class probabilities to assess whether task-specific thresholds improved threshold-dependent metrics such as F1 and balanced accuracy. For regression endpoints, we applied per-task affine calibration of the form ***y_cal_ = ay_pred_ + b*** to correct global prediction bias and scale mismatch without retraining the model. These calibration analyses were used to distinguish representational quality from output calibration effects and to assess whether residual performance gaps were primarily due to imperfect ranking, imperfect threshold choice, or imperfect numerical scaling.

### Model Architecture

#### Overview

Prot2Prop is a parameter-efficient multitask framework for predicting multiple protein developability properties within a single model. The architecture is built around a frozen pretrained ProstT5 encoder, which serves as the shared representation backbone for all downstream tasks (Heinzinger *et al*., 2024). Rather than fully fine-tuning the base language model, Prot2Prop keeps the ProstT5 encoder frozen and trains only lightweight residual modules that operate on the encoder’s outputs, allowing the system to preserve the broad structural and biochemical priors learned during pretraining while keeping the number of trainable parameters modest (Houlsby *et al*., 2019; Schmirler, Heinzinger and Rost, 2024). The model contains a total of 3,185,680 trainable parameters.

#### Input Representation

Each protein is represented as an amino-acid sequence and tokenized using the ProstT5 input format with the **<AA2FOLD>** prefix followed by space-separated residues (Heinzinger *et al.,* 2024). ProstT5 was pretrained as a translation model between amino-acid sequences and Foldseek 3Di structural tokens, a compact alphabet that encodes local backbone conformations (Heinzinger *et al.,* 2024). By providing the **<AA2FOLD>** prefix, the input is processed in the sequence-to-structure direction used during ProstT5 pretraining, allowing the model to leverage structural information learned from large-scale sequence-structure translation. During preprocessing, non-canonical amino acids are normalized to **X** to ensure compatibility with the tokenizer and pretrained backbone. The resulting tokenized sequence is converted into token IDs and passed directly into the frozen ProstT5 encoder to generate residue-level embeddings for downstream multitask prediction.

#### Shared Backbone and Residual Adaptation

Given an input sequence, Prot2Prop first computes token-level embeddings using the frozen ProstT5 encoder (Heinzinger *et al*., 2024). The encoder itself is not updated during training. Instead, the resulting token representations are passed through a **shared residual adapter**, which learns a global multitask correction on top of the pretrained embedding space (Houlsby *et al*., 2019; Schmirler, Heinzinger and Rost, 2024). This adapter consists of layer normalization, a low-dimensional down-projection, GELU activation, an up-projection back to the original embedding dimension, dropout, and a learnable residual scaling parameter. The shared adapter is designed to start near zero, allowing training to introduce task-relevant corrections gradually without disrupting the pretrained backbone.

This shared adapter produces a common token-level representation intended to capture signals that are useful across all developability tasks.

#### Task-Specific Residual Adapters

After shared adaptation, Prot2Prop introduces a second level of specialization through **task-specific residual adapters**. Each prediction task receives its own lightweight adapter applied at the token level (Houlsby *et al*., 2019). This design allows each task to modify the shared representation before pooling, enabling the model to emphasize different residues or local patterns depending on the biochemical endpoint being predicted. For example, the sequence features most relevant to solubility may differ from those most relevant to folding stability or expression yield (Qing *et al*., 2022; Thumuluri *et al*., 2022; Rosace *et al*., 2023).

By applying task-specific adaptation before sequence summarization, Prot2Prop preserves a shared backbone while still allowing each task to reshape the residue-level representation in a targeted way.

#### Attention-Based Sequence Pooling

For each task, the adapted token sequence is converted into a fixed-length sequence representation using **attention pooling** rather than simple mean pooling (Hoang and Singh, 2025). A small feed-forward scoring module assigns learned importance weights across residues, and the final pooled embedding is formed as a weighted combination of token representations. This allows the model to learn which positions contribute most strongly to a given property, rather than treating all residues equally.

Because pooling is performed separately after each task-specific adapter, each task can derive its own sequence summary from a common shared representation.

#### Task Heads

The pooled representation for each task is passed to a dedicated **task-specific prediction head**. Prot2Prop supports both classification and regression tasks in the same model. Classification tasks use a smaller multilayer perceptron head, while regression tasks use a somewhat larger head to provide additional capacity for modeling continuous outputs and correcting scale-sensitive effects. Each head operates independently and produces the final prediction for its associated property.

This design allows Prot2Prop to jointly model heterogeneous endpoints, including binary labels such as solubility or temperature stability and continuous-valued traits such as aggregation propensity, expression yield, and folding stability.

#### Multitask Learning Strategy

Prot2Prop is trained in a multitask setting where each protein may have labels for only a subset of tasks. To support this, the training pipeline uses a **masked-label formulation**: losses are computed only for tasks with observed labels in a given sample. Regression targets are normalized using training-split statistics so that tasks with different value scales contribute more comparably during optimization. Classification tasks use cross-entropy loss, with inverse-frequency class weighting for binary tasks to reduce imbalance effects. The overall multitask loss is computed as the mean of the available per-task losses within each batch.

To reduce domination by tasks with denser label coverage, training batches are sampled using task-aware sample weighting. Sequences contributing labels to rarer tasks receive higher sampling weight, improving balance across objectives during training.

#### Efficiency and Practical Design

The full trainable portion of Prot2Prop consists only of the shared adapter, task-specific residual adapters, attention-pooling module, and task-specific prediction heads, while the underlying ProstT5 encoder remains frozen throughout training (Heinzinger et al., 2024). This design makes Prot2Prop substantially more parameter-efficient than end-to-end fine-tuning of a large protein language model while preserving the ability to learn both shared biochemical structure across tasks and task-specific refinements where needed (Houlsby et al., 2019; Schmirler, Heinzinger and Rost, 2024). By updating only lightweight adaptation modules, Prot2Prop reduces optimization burden, memory requirements, and training complexity, making iterative experimentation with architecture changes, task composition, and calibration strategies substantially more tractable than repeatedly retraining the entire backbone (Houlsby et al., 2019; Schmirler, Heinzinger and Rost, 2024).

A practical advantage of this architecture is that it unifies multiple developability predictions within a single model rather than requiring separate property-specific predictors. The pretrained ProstT5 backbone is loaded once and shared across all downstream tasks, while lightweight task-specific adapters and prediction heads operate on a common embedding space. Compared with conventional single-task workflows, which often duplicate backbone storage and inference for each property, this design reduces redundant computation and improves multi-property throughput (Thumuluri et al., 2022; Pudžiuvelytė et al., 2024). To quantify the practical computational cost of deployment, we compared inference runtime and memory consumption against TemStaPro (Pudžiuvelytė *et al*., 2024) and SaProt (Su *et al*., 2023) (Figure 2).

**Figure 1.**
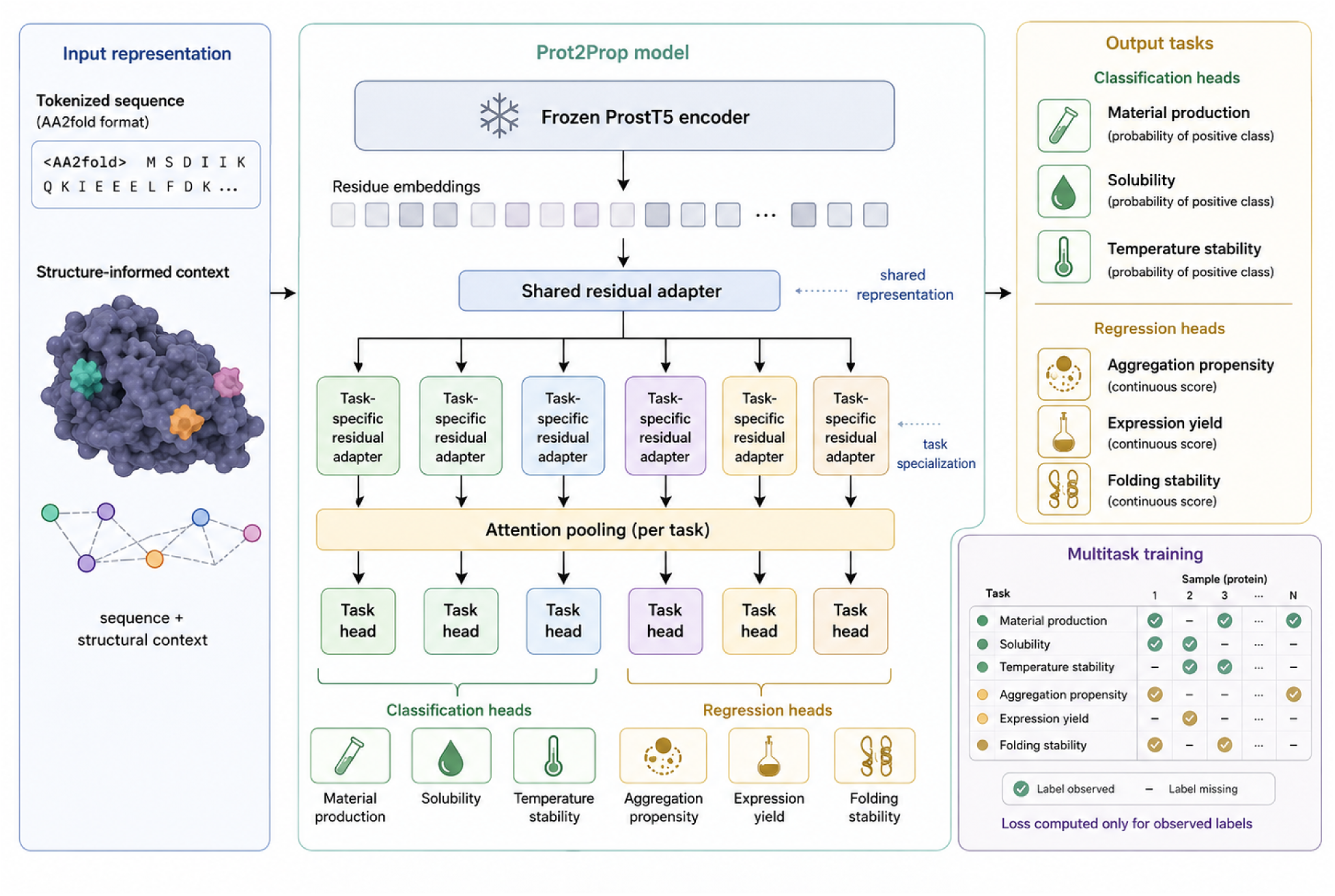
Prot2Prop architecture. A candidate protein sequence is tokenized in AA2fold format and passed through a frozen ProstT5 encoder to generate residue-level embeddings. These embeddings are first processed by a shared residual adapter that captures features common across developability tasks, then routed through task-specific residual adapters that specialize the representation for each endpoint. Per-task attention pooling converts token-level features into fixed-length sequence representations, which are passed to dedicated task heads. Prot2Prop jointly predicts three classification endpoints, material production, solubility, and temperature stability, and three regression endpoints, aggregation propensity, expression yield, and folding stability. During multitask training, losses are computed only for observed labels, allowing proteins with incomplete annotations to contribute to available tasks within a single unified model.

**Figure 2.**
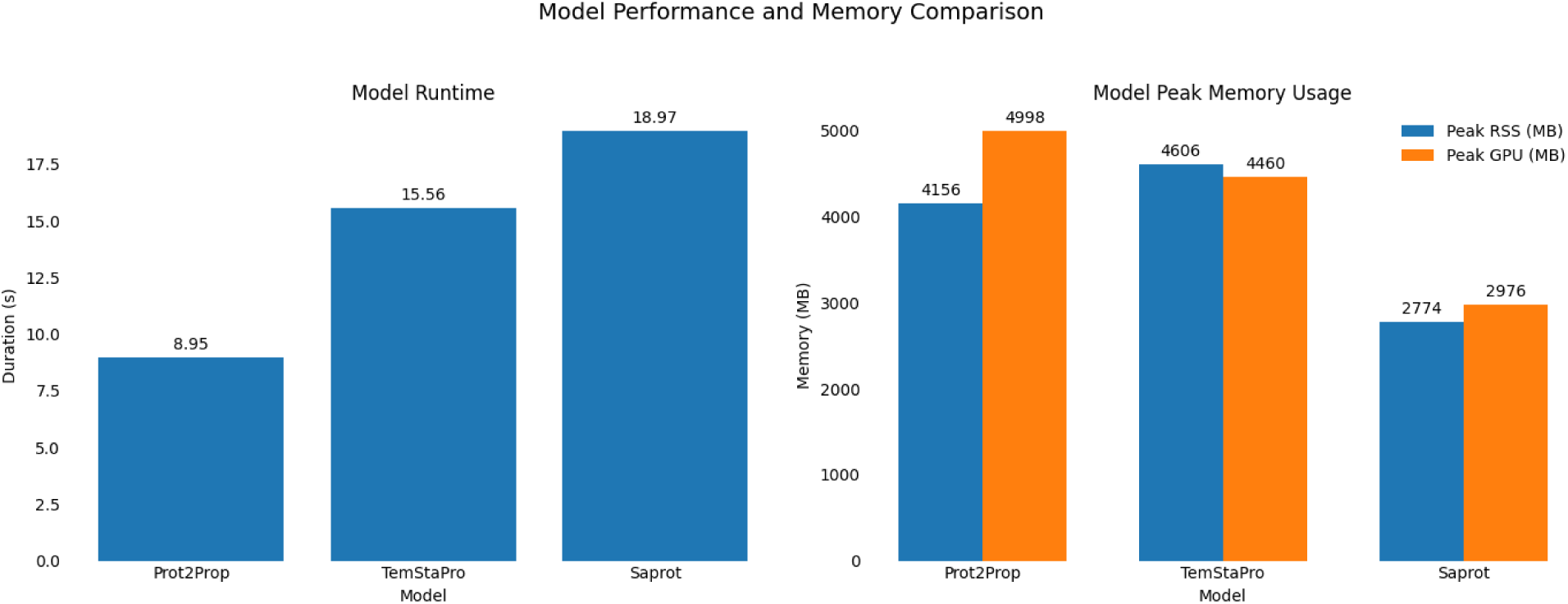
Runtime and memory usage comparison of Prot2Prop, TemStaPro, and SaProt. Runtime (left) and peak memory consumption (right) were measured during inference on identical hardware (Intel Xeon Gold 6342 CPU and NVIDIA A40 GPU). To ensure a fair comparison, benchmarking was performed using the same set of 100 randomly selected sequences from our dataset, each with a maximum length of 72 residues. Prot2Prop achieved the shortest runtime (8.95 s), compared with TemStaPro (15.56 s) and SaProt (18.97 s). Peak CPU memory (RSS) and GPU memory utilization are also shown (Supplementary Table 11). Despite requiring substantially less computational time than the competing methods, Prot2Prop maintained competitive predictive performance, demonstrating a favorable balance between accuracy and computational efficiency. These results highlight the suitability of Prot2Prop for high-throughput developability screening workflows where inference speed and resource utilization are important considerations.

Mixed-precision CUDA execution together with compiler- and optimizer-level acceleration improves hardware utilization during training and inference. A pre-tokenized multitask cache avoids repeated tokenization and enables efficient reuse of sequence lengths and masked labels. Additional batching strategies minimize padding overhead by using length-aware batching, dynamic within-batch padding, and maximum padded-token budgets that prevent unusually long proteins from causing memory spikes. Evaluation batches are sorted by sequence length, while training batches preserve stochasticity through local length-aware sorting. Host-to-device throughput is further improved through pinned-memory loading and asynchronous tensor transfer on CUDA-enabled systems.

A single shared model jointly predicts six developability properties, thereby eliminating the need to train and maintain multiple backbone-level models. Losses are computed only for observed labels, allowing partially annotated proteins to contribute supervision without unnecessary computation on missing targets. Task-aware sample weighting improves coverage of sparsely labeled tasks within a single training run, while early stopping and retention of the best-performing checkpoint reduce wasted computation once validation performance plateaus.

Together, these design choices make Prot2Prop both parameter-efficient and operationally efficient for multi-property protein screening. Under the current setup, a full training run of 10 epochs requires approximately 24 hours on a single NVIDIA L40S GPU, providing a practical turnaround time for multitask experimentation while jointly modeling multiple heterogeneous classification and regression tasks within a unified framework.

## Results

### Iterative model refinement improved multitask performance

Prot2Prop was developed through a series of architectural and training refinements aimed at improving regression quality while preserving strong classification performance across six developability-related tasks: material production, solubility, temperature stability, aggregation propensity, expression yield, and folding stability. The most important changes involved increasing task-head capacity, introducing task-specific residual adapters prior to pooling, testing alternative multitask weighting and ranking-aware objectives, and evaluating optional evolutionary-alignment supervision. Early multitask runs established that a single parameter-efficient model could jointly support heterogeneous classification and regression tasks, but performance on the regression endpoints was initially weaker and less well calibrated. Increasing head capacity in version **2026-04-26** improved overall regression quality and made post-hoc calibration behavior more favorable. In contrast, the ranking-augmented regression-loss variant tested in version **2026-04-28** did not improve the main validation metrics and was not retained. The largest architectural gain came in version **2026-04-29**, which introduced a shared adapter together with task-specific residual adapters applied before pooling. This change substantially improved the regression tasks while leaving classification performance broadly stable, making it the most important design refinement in the development cycle. Learned uncertainty weighting in version **2026-04-30** produced mixed effects, improving some tasks while degrading others, and therefore did not clearly outperform the simpler multitask objective. An additional evolutionary-alignment pathway added in version **2026-05-22** expanded the training framework but did not yet provide a sufficiently strong standalone performance gain to justify treating it as a core contributor to final model quality.

#### Final seeded models and reference checkpoint

After finalizing the architecture, we retrained the model across multiple seeds and selected the strongest checkpoint on validation set performance for use as the primary manuscript reference. Across seeded runs, performance was consistently strong, but the **Seed 1** checkpoint provided the best overall balance across classification and regression tasks (Figure 3).

**Figure 3.**
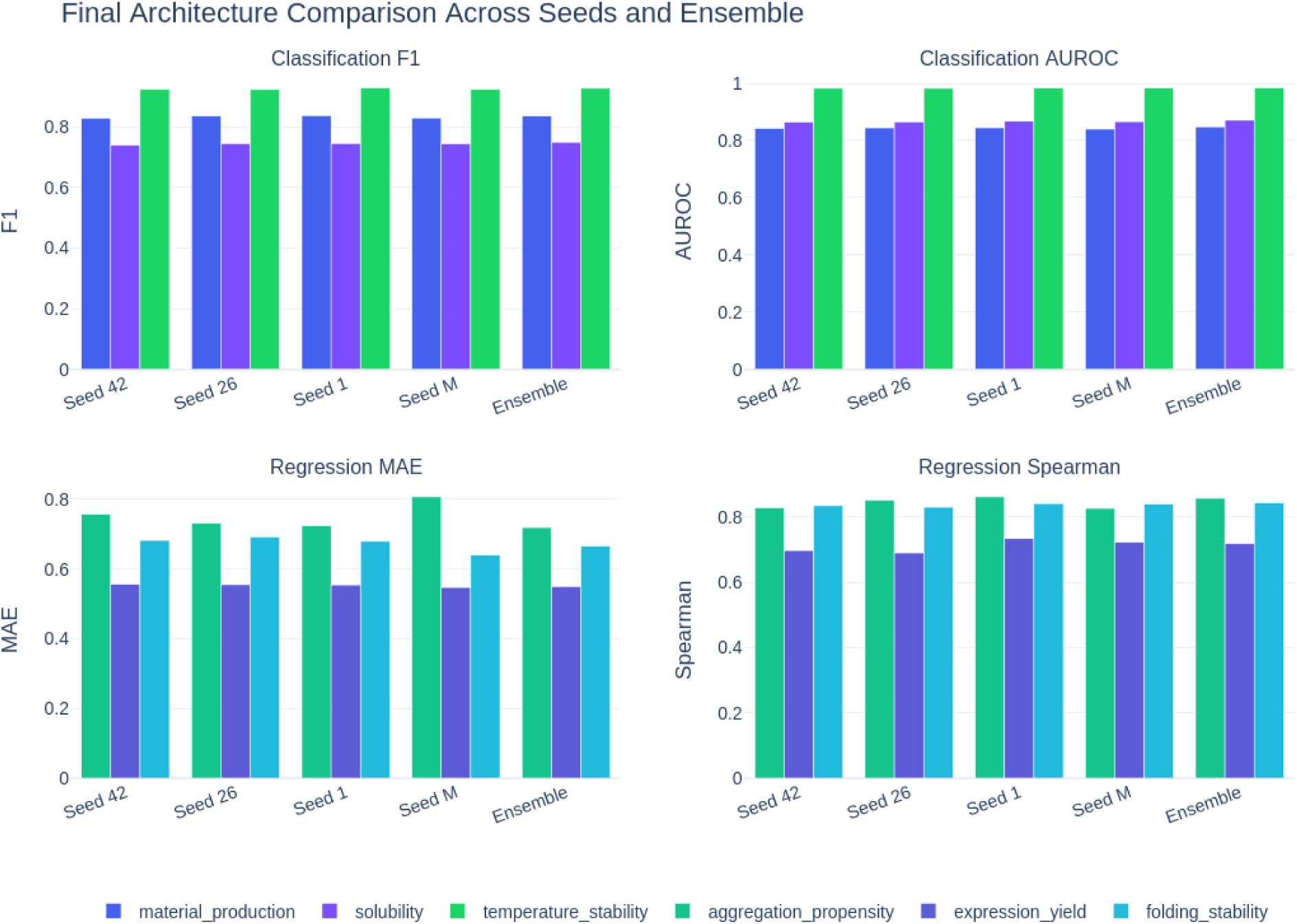
Final architecture performance across seeded checkpoints and ensemble. Performance of the final architecture of Prot2Prop across checkpoints with independent random seeds and the four-checkpoint ensemble (validation set). F1 and AUROC show classification task performance (top), while MAE and Spearman correlation correspond to regression task performance (bottom). Because the classification tasks exhibit varying degrees of class imbalance, we report F1 alongside AUROC to provide a complementary assessment of threshold-dependent classification performance. Performance was largely stable, but regression tasks exhibited greater variation across seeds relative to classification tasks. Seed 1 was chosen as the primary reference checkpoint as it offered the most promising balance across both classification and regression tasks.

On the three classification endpoints, the Seed 1 model achieved **F1 scores of 0.8531, 0.7488, and 0.9308** for material production, solubility, and temperature stability, respectively, with corresponding **AUROC values of 0.8663, 0.8697, and 0.9837**. On the regression tasks, it achieved **MAE/Spearman values of 0.6990/0.8614** for aggregation propensity, **0.5609/0.7321** for expression yield, and **0.6773/0.8384** for folding stability. Post-hoc calibration further improved several endpoints, especially the regression tasks (Figure 4; Table 4). For the Seed 1 checkpoint, aggregation propensity improved from **MAE/RMSE 0.6990/0.9410** to **0.6636/0.8922**, while folding stability improved from **0.6773/0.8791** to **0.4863/0.6405** without changing Spearman correlation. Classification threshold tuning also improved threshold-dependent metrics for some tasks, particularly material production. These results indicate that the learned multitask representation is strong and that a substantial portion of the remaining error on some tasks is calibration-related rather than architectural.

**Figure 4.**
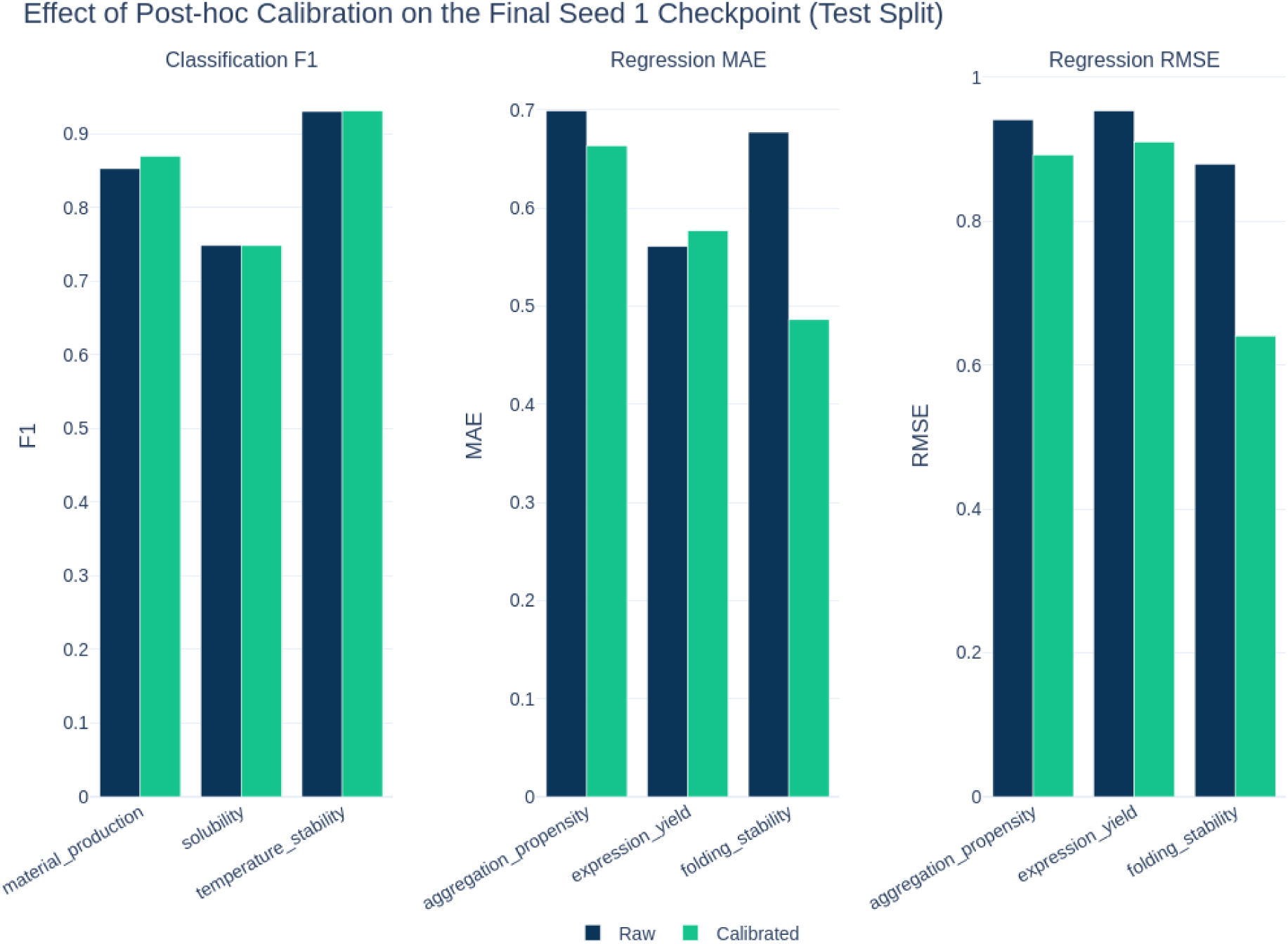
Effect of post-hoc calibration on the final Seed 1 checkpoint. Raw performance compared with the effect of post-hoc calibration for the final checkpoint of Seed 1. Classification task performance is represented by F1 (left), while regression performance is shown using MAE (middle) and RMSE (right). Post-hoc calibration substantially reduced regression error for the aggregation propensity and folding stability tasks, with performance remaining largely stable across the remaining tasks.

#### Summary of architectural findings

Taken together, the Prot2Prop development trajectory supports three main conclusions. First, increasing per-task head capacity beyond minimal linear heads improved regression accuracy and calibration. Second, task-specific token-level adaptation before pooling was the single most effective architectural refinement and produced the clearest overall gain. Third, more complex optimization changes, including ranking-aware regression loss, learned uncertainty weighting, and simple checkpoint ensembling, did not clearly outperform the best single-checkpoint final architecture. Overall, these results support the central claim of this work: a frozen protein language model augmented with lightweight shared and task-specific adapters provides a strong, unified, and computationally practical framework for multitask prediction of protein developability properties.

**Table 1.** Summary of Prot2Prop model development and major architectural changes. Each version is listed with its principal modification, intended purpose, and practical outcome on validation performance. The most consequential improvement came from introducing task-specific residual adapters before pooling in version **2026-04-29**.

| Version | Main change | Intended effect | Outcome |
| --- | --- | --- | --- |
| 2026-04-26 | Replaced minimal linear heads with small MLP heads; removed temperature_stability_abs; added richer validation and calibration analysis | Improve regression expressiveness and calibration | Clear overall improvement |
| 2026-04-28 | Added Huber/ranking-aware regression objective | Improve numeric accuracy while preserving ranking | Slightly worse overall; not retained |
| 2026-04-29 | Added task-specific residual adapters before pooling | Improve task specialization while preserving multitask sharing | Strongest architectural gain |
| 2026-04-30 | Added learned uncertainty weighting | Improve multitask balance | Mixed results |
| 2026-05-22 | Added optional evolutionary-alignment pathway | Add auxiliary supervision for folding stability | Useful infrastructure, limited standalone gain |
| Final seeded runs | Retrained final architecture across seeds | Select best checkpoint | Seed 1 chosen as reference |

**Table 2.** Performance of the final Seed 1 reference checkpoint on classification tasks. Metrics are reported on the held-out test set. Threshold-free metrics are included to separate ranking quality from threshold-dependent classification behavior.

| Task | ACC | Balanced ACC | Precision | Recall | F1 | AUROC | AUPRC |
| --- | --- | --- | --- | --- | --- | --- | --- |
| Material production | 0.7975 | 0.7718 | 0.8726 | 0.8345 | 0.8531 | 0.8663 | 0.9354 |
| Solubility | 0.7730 | 0.7769 | 0.7025 | 0.8015 | 0.7488 | 0.8697 | 0.8358 |
| Temperature stability | 0.9297 | 0.9299 | 0.9074 | 0.9553 | 0.9308 | 0.9837 | 0.9838 |

**Table 3.** Performance of the final Seed 1 reference checkpoint on regression tasks. Regression performance is summarized using absolute error, root mean squared error, and Spearman rank correlation. Reported on the same held-out test set.

| Task | MAE | RMSE | Spearman |
| --- | --- | --- | --- |
| Aggregation propensity | 0.6990 | 0.9410 | 0.8614 |
| Expression yield | 0.5609 | 0.9532 | 0.7321 |
| Folding stability | 0.6773 | 0.8791 | 0.8384 |

**Table 4.** Effect of post-hoc calibration on the final Seed 1 reference checkpoint. Binary tasks were evaluated with tuned decision thresholds, and regression tasks with affine calibration. Calibration produced the largest gains for folding stability and aggregation propensity.

| Task | Raw metric | Calibrated metric |
| --- | --- | --- |
| Material production | F1 0.8531 | F1 0.8699 |
| Solubility | F1 0.7488 | F1 0.7487 |
| Temperature stability | F1 0.9308 | F1 0.9314 |
| Aggregation propensity | MAE/RMSE 0.6990 / 0.9410 | 0.6636 / 0.8922 |
| Expression yield | MAE/RMSE 0.5609 / 0.9532 | 0.5769 / 0.9100 |
| Folding stability | MAE/RMSE 0.6773 / 0.8791 | 0.4863 / 0.6405 |

#### Comparison with a naive one-hot linear baseline

To assess whether Prot2Prop’s performance reflects meaningful gains beyond simple sequence-level encoding, we compared the final validation-selected model against naive one-hot linear baselines. Binary endpoints were evaluated against one-hot logistic regression, and continuous endpoints were evaluated against one-hot ridge regression. This comparison isolates the contribution of the pretrained ProstT5 representation and multitask adapter architecture relative to a simple linear model operating directly on sequence identity.

**Table 5.**
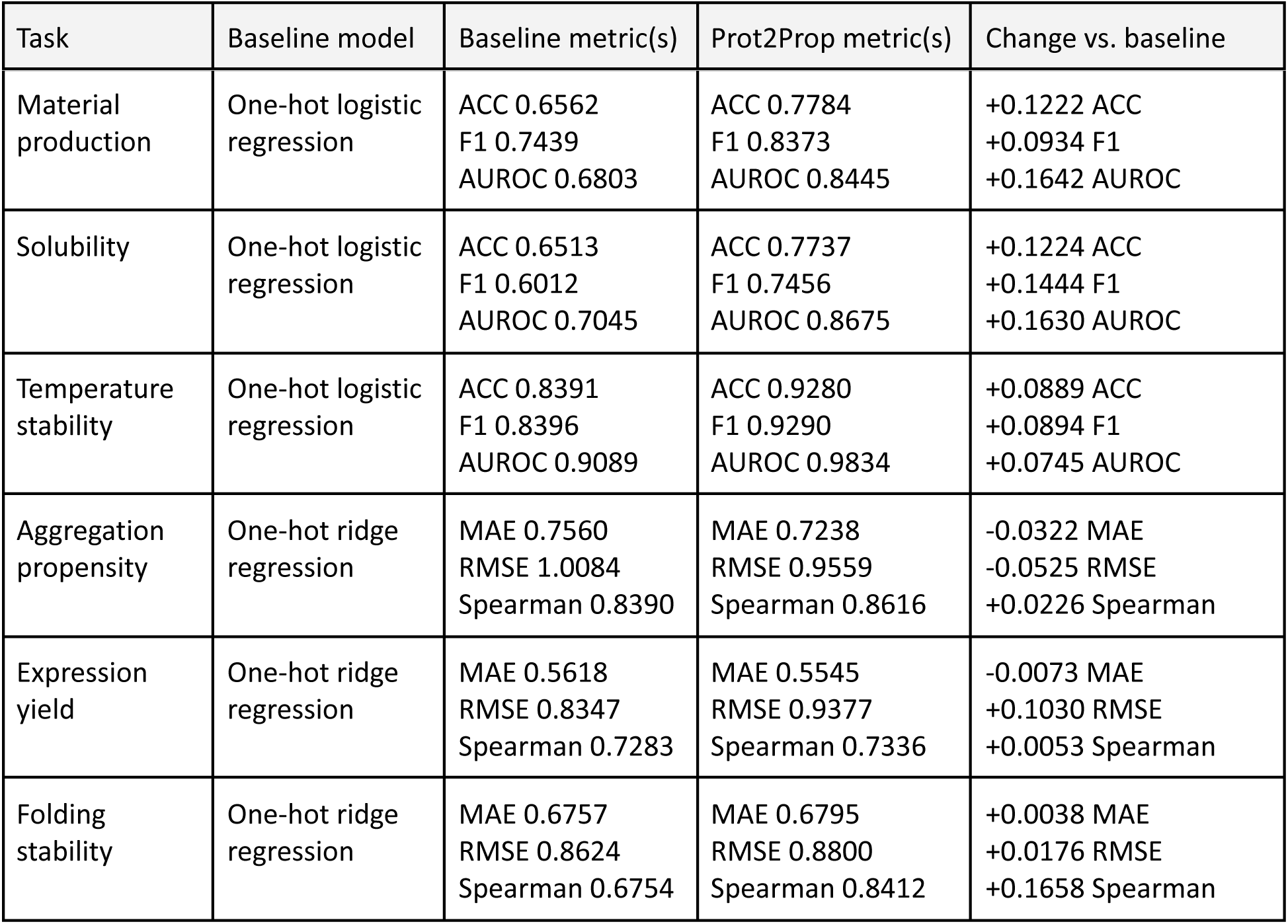
Comparison of Prot2Prop with naive one-hot linear baselines on the held-out validation set. Prot2Prop improved all three classification tasks across accuracy, F1, and AUROC. Regression gains were more task-dependent: aggregation propensity improved across all reported metrics, expression yield improved in MAE and Spearman but not RMSE, and folding stability showed substantially stronger Spearman correlation despite slightly worse raw error metrics.

| Task | Baseline model | Baseline metric(s) | Prot2Prop metric(s) | Change vs. baseline |
| --- | --- | --- | --- | --- |
| Material production | One-hot logistic regression | ACC 0.6562<br>F1 0.7439<br>AUROC 0.6803 | ACC 0.7784<br>F1 0.8373<br>AUROC 0.8445 | +0.1222 ACC<br>+0.0934 F1<br>+0.1642 AUROC |
| Solubility | One-hot logistic regression | ACC 0.6513<br>F1 0.6012<br>AUROC 0.7045 | ACC 0.7737<br>F1 0.7456<br>AUROC 0.8675 | +0.1224 ACC<br>+0.1444 F1<br>+0.1630 AUROC |
| Temperature stability | One-hot logistic regression | ACC 0.8391<br>F1 0.8396<br>AUROC 0.9089 | ACC 0.9280<br>F1 0.9290<br>AUROC 0.9834 | +0.0889 ACC<br>+0.0894 F1<br>+0.0745 AUROC |
| Aggregation propensity | One-hot ridge regression | MAE 0.7560<br>RMSE 1.0084<br>Spearman 0.8390 | MAE 0.7238<br>RMSE 0.9559<br>Spearman 0.8616 | -0.0322 MAE<br>-0.0525 RMSE<br>+0.0226 Spearman |
| Expression yield | One-hot ridge regression | MAE 0.5618<br>RMSE 0.8347<br>Spearman 0.7283 | MAE 0.5545<br>RMSE 0.9377<br>Spearman 0.7336 | -0.0073 MAE<br>+0.1030 RMSE<br>+0.0053 Spearman |
| Folding stability | One-hot ridge regression | MAE 0.6757<br>RMSE 0.8624<br>Spearman 0.6754 | MAE 0.6795<br>RMSE 0.8800<br>Spearman 0.8412 | +0.0038 MAE<br>+0.0176 RMSE<br>+0.1658 Spearman |

Overall, these results indicate that Prot2Prop provides its clearest advantage for classification and rank-based regression performance, while some continuous endpoints remain challenging relative to simple linear baselines.

#### Cross-study comparison with external predictors

To place Prot2Prop performance in context, we compared the calibrated Seed 1 checkpoint with representative external predictors for related protein developability tasks. For classification endpoints, Prot2Prop was compared with NetSolP (Thumuluri *et al*., 2022), SWI (Bhandari, Gardner and Lim, 2020), ESM1b-F (Rives *et al*., 2021), BertThermo (Pei *et al*., 2023), ProLaTherm (Haselbeck *et al*., 2023), and TemStaPro (Pudžiuvelytė *et al*., 2024) where corresponding published metrics were available. For regression endpoints, Prot2Prop folding-stability performance was compared with an external absolute protein folding-stability predictor. Because these external methods were evaluated in their original studies using different datasets, label definitions, temperature thresholds, target units, and metric sets, the comparisons should be interpreted as cross-study context rather than as a controlled head-to-head benchmark. Values are reported exactly as available from the source publications, with unreported metrics marked as NR and correlation coefficients kept separate from Spearman rank correlations when the correlation type was not explicitly specified.

**Table 6.** Cross-study comparison of Prot2Prop classification performance against task-specific protein property predictors. This table compares the Prot2Prop Seed 1 checkpoint against representative external models for related binary protein developability tasks. Comparator values are taken directly from the original publications and are reported only for metrics explicitly provided by those studies. NetSolP, SWI, and ESM1b-F are included for solubility and material-production-related usability because they report performance on the Price solubility and usability benchmarks, where usability reflects a combined soluble-expression criterion. TemStaPro, BertThermo, and ProLaTherm are included as the external comparators for temperature stability. Because the temperature-stability comparators report performance at multiple temperature thresholds, their metrics are shown as ranges representing the minimum and maximum values observed among evaluations using the 40°C, 45°C, 50°C, 55°C, 60°C, and 65°C datasets/settings, rather than as single point estimates. Metrics that were not reported in the corresponding source are marked as NR. This comparison is intended to contextualize Prot2Prop relative to established task-specific predictors, while recognizing that the benchmark datasets, label definitions, and evaluation protocols differ across studies. *Prot2Prop values are from version **2026-05-22**, Seed 1 checkpoint.

| Model Evaluated | Task Evaluated | Accuracy | Precision | AUROC |
| --- | --- | --- | --- | --- |
| Prot2Prop* | material_production | **0.7975** | **0.8726** | **0.8663** |
| NetSolP | material_production / usability | 0.65 ± 0.02 | 0.65 ± 0.04 | 0.70 ± 0.01 |
| ESM1b-F | material_production / usability | 0.65 ± 0.01 | 0.64 ± 0.03 | 0.71 ± 0.01 |
| SWI | material_production / usability | 0.58 ± 0.01 | 0.54 ± 0.01 | 0.64 ± 0.02 |
| Prot2Prop* | solubility | **0.7730** | 0.7025 | **0.8697** |
| NetSolP | solubility | 0.728 | **0.773** | 0.760 |
| ESM1b-F | solubility | 0.723 | 0.764 | 0.761 |
| SWI | solubility | 0.680 | 0.712 | 0.690 |
| Prot2Prop* | temperature_stability | **0.9297** | **0.9074** | **0.9837** |
| TemStaPro | temperature_stability | 0.817-0.919 | 0.449-0.818 | 0.908-0.978 |
| BertThermo | temperature_stability | 0.683-0.811 | 0.767-0.792 | 0.683-0.811 |
| ProLaTherm | temperature_stability | 0.679-0.866 | 0.886-0.911 | 0.680-0.865 |

For regression endpoints, Prot2Prop was compared with SaProtΔG and ESM3ΔG for folding stability (Cho *et al*., 2026); no external predictor with compatible evaluation metrics was identified for aggregation propensity or expression yield (Table 7).

**Table 7.** Cross-study comparison of Prot2Prop regression performance against task-specific protein property predictors. Prot2Prop values are reported from the Seed 1 checkpoint before (raw) and after (calibrated) post-hoc affine calibration. Calibrated Prot2Prop values correspond to MAE and RMSE from Table 4; Spearman correlation is unchanged by affine calibration and is taken from Table 3. No external comparator with compatible evaluation metrics was identified for aggregation propensity or expression yield; only folding stability can be contextualized against published regression benchmarks. Because benchmark datasets, label definitions, target units, and evaluation protocols differ across studies, values from external sources should be interpreted as cross-study context rather than a controlled head-to-head comparison. Metrics not reported in the corresponding source are marked as NR. *Prot2Prop values are from version **2026-05-22**, Seed 1 checkpoint. †SaProtΔG and ESM3ΔG values are taken from Cho et al. (2026) and are reported on the MGnify Stability held-out test set. These results are included only as cross-study context rather than as a controlled head-to-head benchmark, because the datasets, target definitions, measurement scales, and evaluation protocols differ from those used for Prot2Prop. In particular, the Prot2Prop folding stability task is modeled as a ProteinGym-derived relative score rather than an absolute ΔG prediction. MAE was not reported in the source publication (NR). Spearman correlation is shown because it is unit-free and is therefore the most interpretable metric across studies with differing target scales.

| Model Evaluated | Task Evaluated | MAE | RMSE | Spearman |
| --- | --- | --- | --- | --- |
| Prot2Prop* (raw) | aggregation propensity | 0.6990 | 0.9410 | 0.8614 |
| Prot2Prop* (calibrated) | aggregation propensity | **0.6636** | **0.8922** | **0.8614** |
| Prot2Prop* (raw) | expression yield | **0.5609** | 0.9532 | 0.7321 |
| Prot2Prop* (calibrated) | expression yield | 0.5769 | **0.9100** | 0.7321 |
| Prot2Prop* (raw) | folding stability | 0.6773 | 0.8791 | 0.8384 |
| Prot2Prop* (calibrated) | folding stability | **0.4863** | **0.6405** | 0.8384 |
| SaProt$\Delta G$† | folding stability ($\Delta G$) | NR | 0.80 | **0.88** |
| ESM3$\Delta G$† | folding stability ($\Delta G$) | NR | 0.80 | 0.87 |

#### Supplementary Results: version-by-version performance analysis

To document the development trajectory of Prot2Prop in full, we evaluated each major architectural version across the same six-task multitask benchmark. The earliest strong baseline, version **2026-04-26**, demonstrated that increasing task-head capacity beyond a minimal linear projection substantially improved regression behavior and made post-hoc calibration more effective. In particular, this version established a stronger baseline for folding stability and expression yield, while also introducing richer classification reporting through AUROC and AUPRC. Version **2026-04-28** tested whether a ranking-aware regression objective could preserve or improve Spearman correlation while also strengthening numeric prediction. In practice, this approach slightly degraded both classification and regression metrics and did not justify the added objective complexity. Version **2026-04-29** introduced the key architectural change of shared adaptation followed by task-specific residual token-level adaptation before pooling. This produced the strongest improvement in the entire development cycle, with clear gains in regression MAE, RMSE, and Spearman correlation across aggregation propensity, expression yield, and folding stability. Classification performance remained stable, indicating that the added specialization primarily benefited the harder continuous-valued tasks without compromising the binary endpoints. Version **2026-04-30** added learned uncertainty weighting to rebalance multitask optimization. Although some tasks improved, especially aggregation propensity and temperature stability, other tasks regressed, leaving the net effect mixed. Version **2026-05-22** then added an optional evolutionary-alignment pathway intended mainly for folding stability. This extension broadened the training framework but did not yet show a decisive top-line performance gain sufficient to justify making it a core part of the main model narrative. After selecting the final architecture, we retrained across multiple seeds. These runs showed that performance was stable overall but still exhibited meaningful seed-to-seed variation, particularly on the regression tasks. Among all final checkpoints, Seed 1 provided the strongest overall balance and was therefore selected as the main manuscript reference (Supplementary Figure 1).

## Discussion and Conclusions

Prot2Prop was motivated by a practical limitation in current protein developability modeling: most available approaches treat each property as an isolated prediction problem, even though many relevant traits are biologically coupled and often need to be assessed together in realistic engineering workflows. A multitask framework is therefore attractive for both biological and operational reasons. Shared training can capture common developability-related signal across endpoints, while a single unified model avoids the inefficiency of maintaining and running separate predictors for each property. This motivation is consistent with prior work showing the value of multitask learning in biological prediction (Ramsundar *et al*., 2015) and protein language model adaptation (An and Weng, 2022; Schmirler, Heinzinger and Rost, 2024).

The present results support this design choice. Across six classification and regression endpoints, Prot2Prop achieved strong overall performance while remaining parameter-efficient through a frozen ProstT5 backbone and lightweight adaptation modules. The most important architectural gain came from introducing task-specific residual adapters before pooling, which improved regression performance without materially harming classification. This pattern suggests that the main challenge was not simply insufficient shared representation capacity, but the need for task-dependent token reshaping and task-specific sequence summarization. In that sense, Prot2Prop appears to benefit from combining shared multitask transfer with explicit task-level specialization rather than relying on a single common sequence representation for all endpoints.

The comparison with the naive one-hot linear baseline helps clarify where these gains arise. Prot2Prop clearly outperformed the baseline across all three classification tasks and showed stronger rank-based performance on all regression endpoints, especially folding stability. At the same time, some regression metrics remained closer to the linear baseline, indicating that not every developability endpoint benefits equally from richer contextual representations. This is an important result rather than a negative one: it suggests that some properties may have a larger linear or composition-driven component, whereas others depend more strongly on higher-order contextual signal captured by pretrained protein language models.

Several limitations should also be acknowledged. First, the comparisons to external predictors are cross-study rather than direct head-to-head evaluations on identical datasets and splits, and should therefore be interpreted cautiously. Second, the current study does not yet include matched single-task baselines using the same backbone and preprocessing pipeline, which limits how precisely the gains can be attributed to multitask transfer itself. Third, the present task set, while practically relevant, still covers only a subset of developability space and does not yet address more context-dependent properties such as partner-specific binding affinity, substrate-conditioned activity, or host-specific expression. Even with these limitations, the current results show that parameter-efficient multitask adaptation (Houlsby *et al*., 2019; Sledzieski *et al*., 2024) of a protein language model (Heinzinger *et al*., 2024) can provide a strong, extensible, and computationally practical framework for unified developability prediction.

## Future Directions

Several natural extensions follow from this work. A first priority is direct head-to-head evaluation against representative baseline models on identical data splits and protocols, together with matched single-task baselines, so that the benefits of multitask learning can be isolated more cleanly. A second priority is broader backbone comparison, including other protein language models and alternative parameter-efficient adaptation strategies such as LoRA or qLoRA. At present, however, it is not obvious that a shared-backbone LoRA approach would outperform the current design, because the strongest gains so far appear to come from larger task heads and task-specific residual adapters before pooling rather than from simply increasing backbone flexibility.

A third direction is expansion of both task scope and biological context. Immediate extensions include additional developability-related endpoints such as pH or ionic-strength stability, secretion signal prediction, oligomerization propensity, metal-binding or cofactor dependence, immunogenicity, and epitope likelihood. Longer-term extensions could incorporate richer contextual inputs to support tasks such as binding affinity, enzymatic activity, host-specific expression, localization, and post-translational modification propensity. Expansion into antibody and biologic developability is especially promising, where multitask modeling of solubility, stability, aggregation risk, and immunogenicity could be directly relevant for therapeutic design. Related recent work in multitask (Pourmirzaei *et al*., 2024; H. Wang *et al*., 2025), multi-objective (Alsamkary *et al*., 2025), and meta-learning-based biological modeling (Zhou *et al*., 2024; Beck *et al*., 2025) suggests that these directions are well motivated and potentially high value.

## Data & Code Availability

All source code for training, validation, and inference, together with the trained Prot2Prop model weights, is available at https://github.com/NeurosnapInc/Prot2Prop/ under the Apache License 2.0.

A public web server for running Prot2Prop is available through the Neurosnap platform at: https://neurosnap.ai/service/Prot2Prop

## Acknowledgements

We thank Dr. Anthony Gitter for reviewing the manuscript, providing valuable feedback and suggestions, and participating in discussions throughout the development of this work.

## Author Contributions

K.A. conceived and supervised the project and led the design and training of the models. D.G.A. contributed to model design and training and assisted in the preparation and writing of the manuscript. D.S.K. and C.J. contributed to data curation and dataset preparation. M.S. provided feedback throughout the development of this project and assisted in the review and preparation of the manuscript.

## Ethics Declarations

Authors K.A. and D.G.A. are affiliated with Neurosnap Inc., which may benefit financially from the development and validation of Prot2Prop as part of its suite of bioinformatics services.

## Supplementary Information

This supplementary section provides additional data, figures, and methodological details supporting the main findings of the Prot2Prop study. It includes in-depth definitions and visual distributions of all computational metrics used to assess developability prediction performance, along with version-by-version performance analyses, post-hoc calibration effects, and comparative analyses across models and modalities.

## Development and Performance Tables

**Supplementary Table 1.** Headline validation metrics for Prot2Prop version 2026-04-26. This version introduced higher-capacity task heads and richer evaluation metrics, resulting in clear overall improvement relative to earlier baselines.

| Task | Metric 1 | Metric 2 | Metric 3 |
| --- | --- | --- | --- |
| Material production | F1 0.8393 | AUROC 0.8463 | AUPRC 0.9265 |
| Solubility | F1 0.7397 | AUROC 0.8619 | AUPRC 0.8231 |
| Temperature stability | F1 0.9234 | AUROC 0.9829 | AUPRC 0.9832 |
| Aggregation propensity | MAE 0.8434 | RMSE 1.1004 | Spearman 0.7860 |
| Expression yield | MAE 0.6129 | RMSE 1.0296 | Spearman 0.6740 |
| Folding stability | MAE 0.7812 | RMSE 0.9900 | Spearman 0.8133 |

**Supplementary Table 2.** Headline validation metrics for Prot2Prop version 2026-04-28. Adding a ranking-aware regression objective did not improve overall performance and was not retained.

| Task | Metric 1 | Metric 2 | Metric 3 |
| --- | --- | --- | --- |
| Material production | F1 0.8349 | AUROC 0.8459 | AUPRC 0.9259 |
| Solubility | F1 0.7380 | AUROC 0.8592 | AUPRC 0.8190 |
| Temperature stability | F1 0.9170 | AUROC 0.9837 | AUPRC 0.9839 |
| Aggregation propensity | MAE 0.8551 | RMSE 1.1350 | Spearman 0.7781 |
| Expression yield | MAE 0.6418 | RMSE 1.0886 | Spearman 0.6688 |
| Folding stability | MAE 0.8535 | RMSE 1.0647 | Spearman 0.7964 |

**Supplementary Table 3.** Headline validation metrics for Prot2Prop version 2026-04-29. Task-specific residual adapters before pooling produced the strongest regression improvement while preserving classification performance.

| Task | Metric 1 | Metric 2 | Metric 3 |
| --- | --- | --- | --- |
| Material production | F1 0.8362 | AUROC 0.8458 | AUPRC 0.9262 |
| Solubility | F1 0.7443 | AUROC 0.8640 | AUPRC 0.8296 |
| Temperature stability | F1 0.9233 | AUROC 0.9838 | AUPRC 0.9842 |
| Aggregation propensity | MAE 0.7268 | RMSE 0.9547 | Spearman 0.8452 |
| Expression yield | MAE 0.5464 | RMSE 0.9304 | Spearman 0.7267 |
| Folding stability | MAE 0.6260 | RMSE 0.8305 | Spearman 0.8373 |

**Supplementary Table 4.** Headline validation metrics for Prot2Prop version 2026-04-30. Learned uncertainty weighting changed the multitask tradeoff but did not clearly improve net performance.

| Task | Metric 1 | Metric 2 | Metric 3 |
| --- | --- | --- | --- |
| Material production | F1 0.8359 | AUROC 0.8437 | AUPRC 0.9247 |
| Solubility | F1 0.7424 | AUROC 0.8660 | AUPRC 0.8300 |
| Temperature stability | F1 0.9287 | AUROC 0.9840 | AUPRC 0.9842 |
| Aggregation propensity | MAE 0.6876 | RMSE 0.9205 | Spearman 0.8522 |
| Expression yield | MAE 0.5564 | RMSE 0.9491 | Spearman 0.7117 |
| Folding stability | MAE 0.6883 | RMSE 0.8927 | Spearman 0.8322 |

**Supplementary Table 5.** Headline validation metrics for Prot2Prop version 2026-05-22. The optional evolutionary-alignment pathway expanded training support but did not produce a decisive standalone performance gain.

| Task | Metric 1 | Metric 2 | Metric 3 |
| --- | --- | --- | --- |
| Material production | F1 0.8338 | AUROC 0.8408 | AUPRC 0.9222 |
| Solubility | F1 0.7455 | AUROC 0.8667 | AUPRC 0.8315 |
| Temperature stability | F1 0.9257 | AUROC 0.9836 | AUPRC 0.9839 |
| Aggregation propensity | MAE 0.7264 | RMSE 0.9615 | Spearman 0.8550 |
| Expression yield | MAE 0.5473 | RMSE 0.9305 | Spearman 0.7289 |
| Folding stability | MAE 0.6435 | RMSE 0.8410 | Spearman 0.8440 |

**Supplementary Table 6.** Performance of the final Seed 1 reference checkpoint on classification tasks. Metrics are reported on the held-out validation set. Threshold-free metrics are included to separate ranking quality from threshold-dependent classification behavior.

| Task | ACC | Balanced ACC | Precision | Recall | F1 | AUROC | AUPRC |
| --- | --- | --- | --- | --- | --- | --- | --- |
| Material production | 0.7784 | 0.7565 | 0.8662 | 0.8103 | 0.8373 | 0.8445 | 0.9268 |
| Solubility | 0.7737 | 0.7761 | 0.7053 | 0.7909 | 0.7456 | 0.8675 | 0.8306 |
| Temperature stability | 0.9280 | 0.9282 | 0.9076 | 0.9515 | 0.9290 | 0.9834 | 0.9837 |

**Supplementary Table 7.** Performance of the final Seed 1 reference checkpoint on regression tasks. Regression performance is summarized using absolute error, root mean squared error, and Spearman rank correlation.

| Task | MAE | RMSE | Spearman |
| --- | --- | --- | --- |
| Aggregation propensity | 0.7238 | 0.9559 | 0.8616 |
| Expression yield | 0.5545 | 0.9377 | 0.7336 |
| Folding stability | 0.6795 | 0.8800 | 0.8412 |

**Supplementary Table 8.**
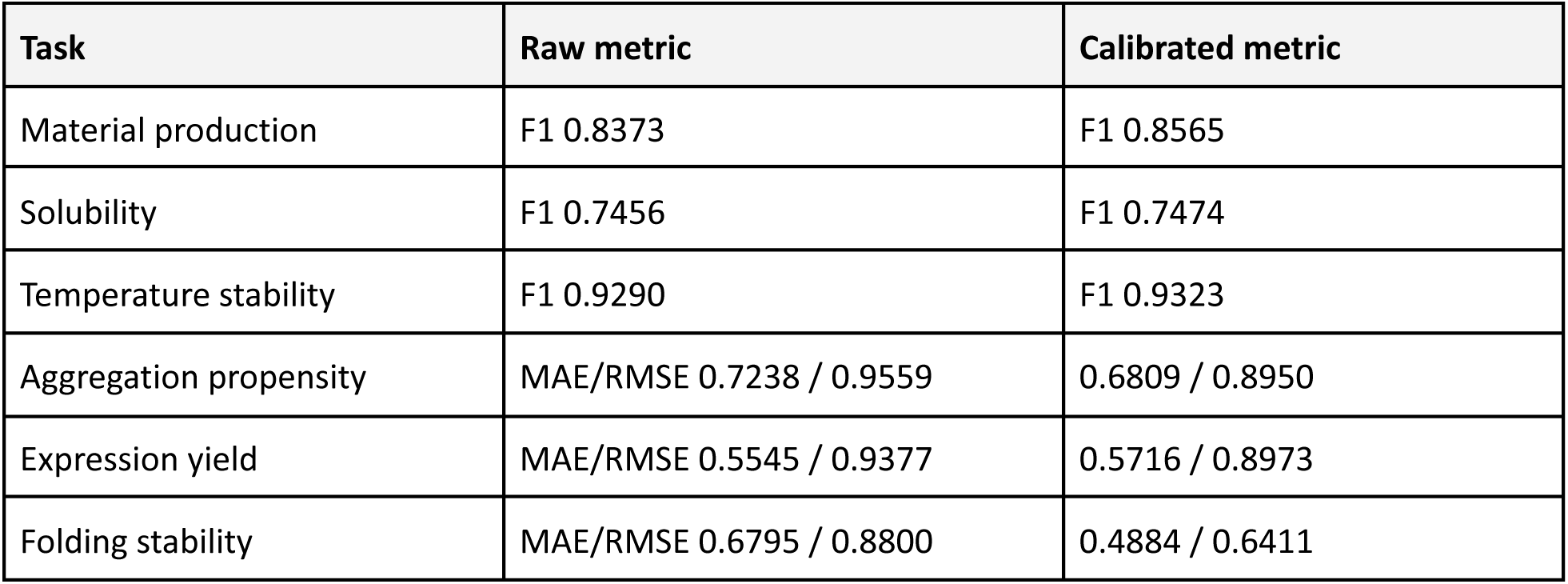
Effect of post-hoc calibration on the final Seed 1 reference checkpoint. Binary tasks were evaluated with tuned decision thresholds, and regression tasks with affine calibration. Calibration produced the largest gains for folding stability and aggregation propensity.

| Task | Raw metric | Calibrated metric |
| --- | --- | --- |
| Material production | F1 0.8373 | F1 0.8565 |
| Solubility | F1 0.7456 | F1 0.7474 |
| Temperature stability | F1 0.9290 | F1 0.9323 |
| Aggregation propensity | MAE/RMSE 0.7238 / 0.9559 | 0.6809 / 0.8950 |
| Expression yield | MAE/RMSE 0.5545 / 0.9377 | 0.5716 / 0.8973 |
| Folding stability | MAE/RMSE 0.6795 / 0.8800 | 0.4884 / 0.6411 |

**Supplementary Table 9.**
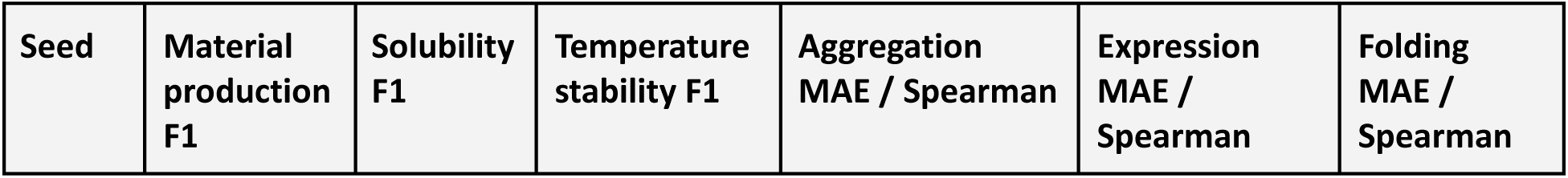
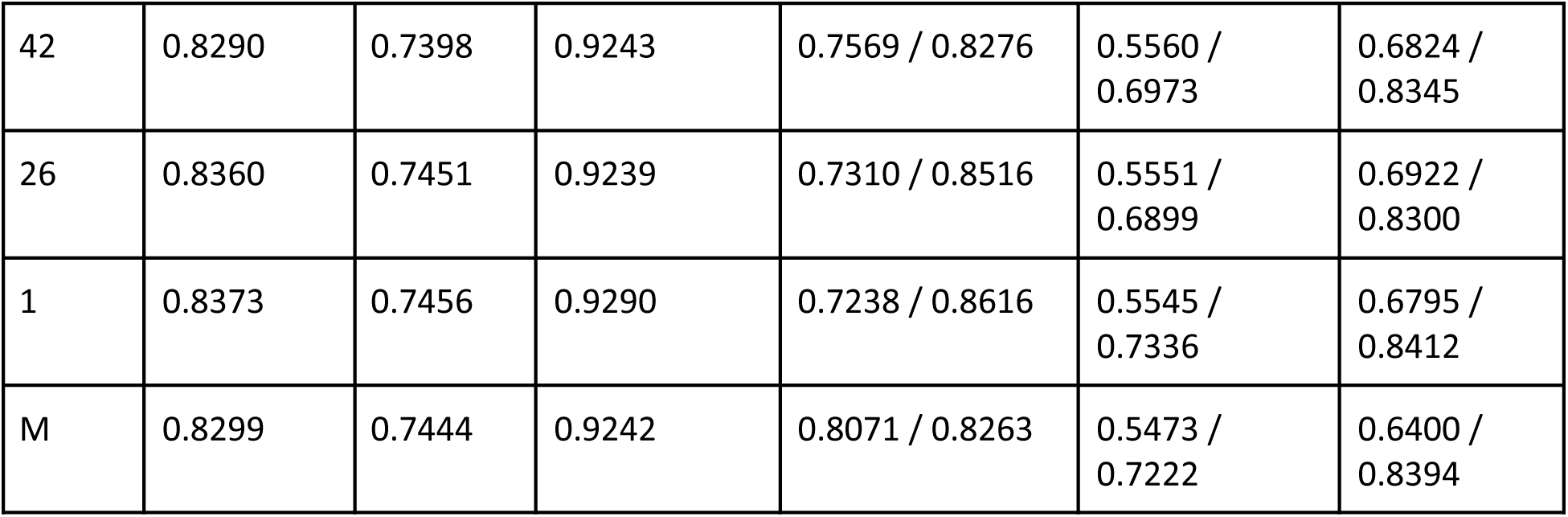
Seed-to-seed comparison for the final Prot2Prop architecture. Seed 1 provided the strongest overall balance across classification and regression and was selected as the manuscript reference checkpoint.

| Seed | Material production F1 | Solubility F1 | Temperature stability F1 | Aggregation MAE / Spearman | Expression MAE / Spearman | Folding MAE / Spearman |
| --- | --- | --- | --- | --- | --- | --- |
| 42 | 0.8290 | 0.7398 | 0.9243 | 0.7569 / 0.8276 | 0.5560 / 0.6973 | 0.6824 / 0.8345 |
| 26 | 0.8360 | 0.7451 | 0.9239 | 0.7310 / 0.8516 | 0.5551 / 0.6899 | 0.6922 / 0.8300 |
| 1 | 0.8373 | 0.7456 | 0.9290 | 0.7238 / 0.8616 | 0.5545 / 0.7336 | 0.6795 / 0.8412 |
| M | 0.8299 | 0.7444 | 0.9242 | 0.8071 / 0.8263 | 0.5473 / 0.7222 | 0.6400 / 0.8394 |

**Supplementary Table 10.** Headline performance of the four-checkpoint ensemble. Although the ensemble was competitive, it did not consistently outperform the best single-checkpoint reference model.

| Task | Metric 1 | Metric 2 | Metric 3 |
| --- | --- | --- | --- |
| Material production | F1 0.8360 | AUROC 0.8480 | AUPRC 0.9283 |
| Solubility | F1 0.7499 | AUROC 0.8714 | AUPRC 0.8369 |
| Temperature stability | F1 0.9279 | AUROC 0.9840 | AUPRC 0.9844 |
| Aggregation propensity | MAE 0.7190 | RMSE 0.9475 | Spearman 0.8575 |
| Expression yield | MAE 0.5498 | RMSE 0.9393 | Spearman 0.7182 |
| Folding stability | MAE 0.6660 | RMSE 0.8661 | Spearman 0.8432 |

**Supplementary Table 11.**
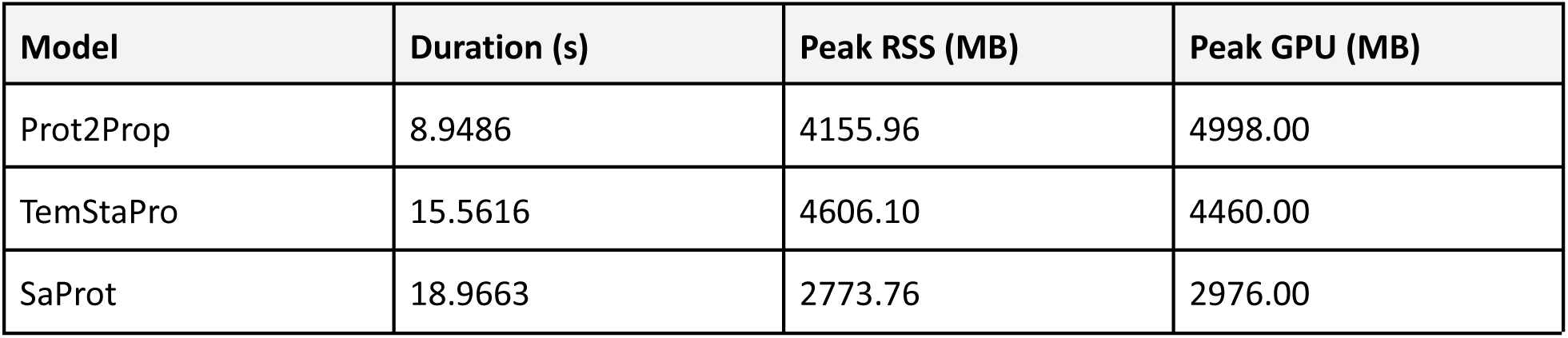
Comparison of computational resource usage and duration for Prot2Prop and external models. Benchmarking was performed on an Intel® Xeon® Gold 6342 CPU running at 2.80 GHz and an NVIDIA A40 GPU. The system used NVIDIA driver version 570.211.01 with CUDA 12.8. The GPU power limit was set to 300 W, and the device operated in P8 power state during testing.

**Supplementary Figure 1.**
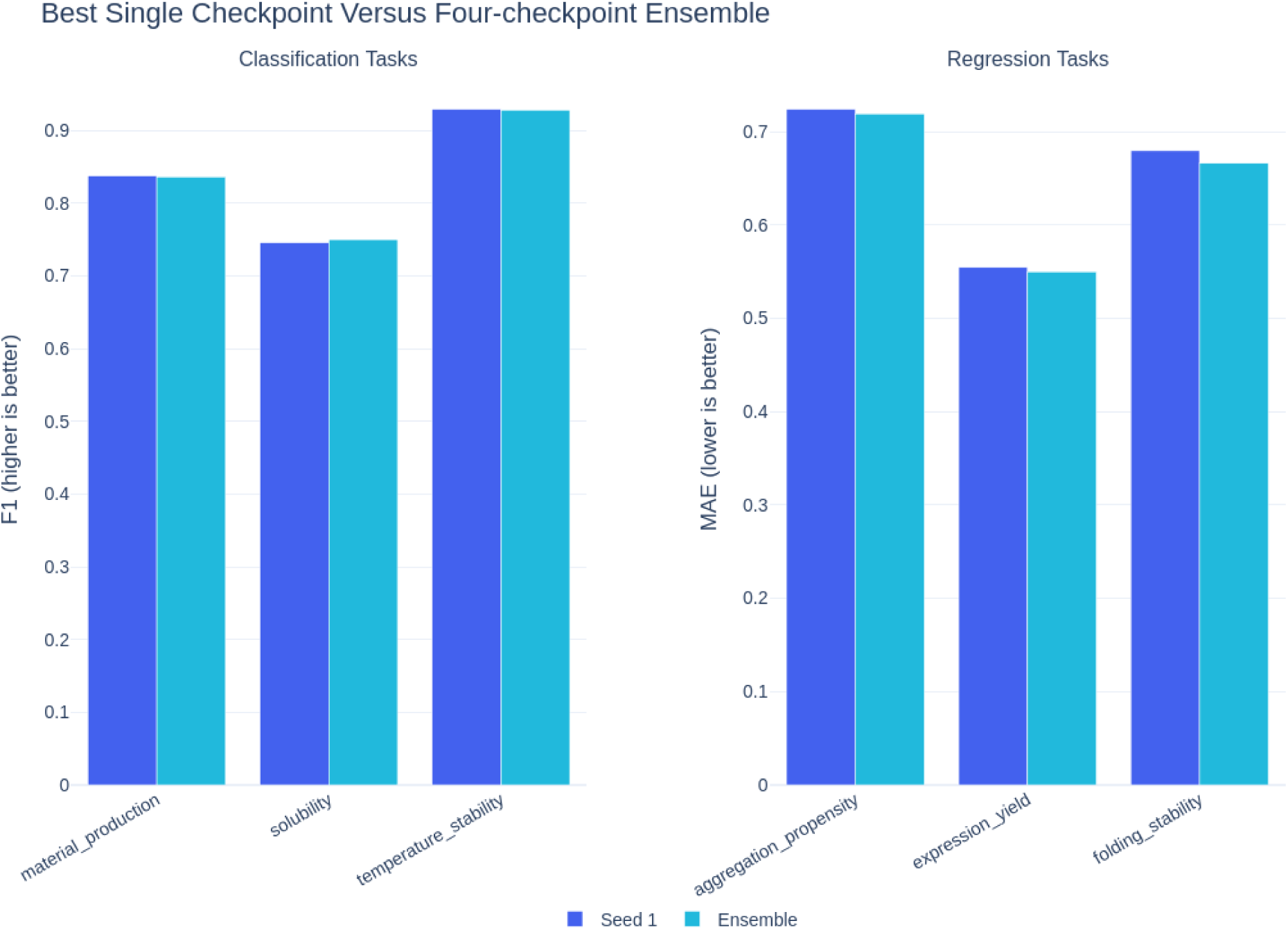
Best single checkpoint versus four-checkpoint ensemble. Performance comparison of the best final checkpoint and the four-checkpoint ensemble (validation set). Classification performance is showcased by F1 (left), while regression task performance is measured with MAE (right). There was no obvious advantage conferred by the four-checkpoint ensemble approach, which supported Seed 1’s status as the primary manuscript reference checkpoint.

## References

Alsamkary, H. et al. (2025) “Ankh3: Multi-Task Pretraining with Sequence Denoising and Completion Enhances Protein Representations.” arXiv. Available at: 10.48550/arXiv.2505.20052.

An, J. and Weng, X. (2022) “Collectively encoding protein properties enriches protein language models,” BMC Bioinformatics, 23(1), p. 467. Available at: 10.1186/s12859-022-05031-z.

Anyaegbunam, U.A. et al. (2026) “Cross-Domain Transfer Learning from Peptides to Lipids Using a Multi-Property Fine-Tuned LLM.” Bioinformatics. Available at: 10.64898/2026.01.06.697904.

Beck, J. et al. (2025) “Metalic: Meta-Learning In-Context with Protein Language Models.” arXiv. Available at: 10.48550/arXiv.2410.08355.

Bhandari, B.K., Gardner, P.P. and Lim, C.S. (2020) “Solubility-Weighted Index: fast and accurate prediction of protein solubility,” Bioinformatics. Edited by Y. Ponty, 36(18), pp. 4691–4698. Available at: 10.1093/bioinformatics/btaa578.

Brandes, N., et al. (2022) “ProteinBERT: a universal deep-learning model of protein sequence and function,” Bioinformatics. Edited by P.L. Martelli, 38(8), pp. 2102–2110. Available at: 10.1093/bioinformatics/btac020.

Chandra, A. et al. (2023) “Transformer-based deep learning for predicting protein properties in the life sciences,” eLife, 12, p. e82819. Available at: 10.7554/eLife.82819.

Cho, Y. et al. (2026) “Accurate protein stability prediction for small domains using mega-scale experiments.” Biophysics. Available at: 10.64898/2026.05.19.726285.

Haselbeck, F. et al. (2023) “Superior protein thermophilicity prediction with protein language model embeddings,” NAR Genomics and Bioinformatics, 5(4), p. lqad087. Available at: 10.1093/nargab/lqad087.

Heinzinger, M. et al. (2024) “Bilingual language model for protein sequence and structure,” NAR Genomics and Bioinformatics, 6(4), p. lqae150. Available at: 10.1093/nargab/lqae150.

Hoang, M. and Singh, M. (2025) “Locality-aware pooling enhances protein language model performance across varied applications,” Bioinformatics, 41(Supplement_1), pp. i217–i226. Available at: 10.1093/bioinformatics/btaf178.

Houlsby, N., et al. (2019) “Parameter-Efficient Transfer Learning for NLP,” in K. Chaudhuri and R. Salakhutdinov (eds.) Proceedings of the 36th International Conference on Machine Learning. PMLR (Proceedings of Machine Learning Research), pp. 2790–2799. Available at: https://proceedings.mlr.press/v97/houlsby19a.html.

Kuriata, A. et al. (2019) “Aggrescan3D (A3D) 2.0: prediction and engineering of protein solubility,” Nucleic Acids Research, 47(W1), pp. W300–W307. Available at: 10.1093/nar/gkz321.

Lin, Z. et al. (2023) “Evolutionary-scale prediction of atomic-level protein structure with a language model,” Science, 379(6637), pp. 1123–1130. Available at: 10.1126/science.ade2574.

Neurosnap Inc. (2022) “Neurosnap: An Online Platform for Computational Biology and Chemistry.” Available at: https://neurosnap.ai/.

Notin, P. et al. (2023) “ProteinGym: Large-Scale Benchmarks for Protein Design and Fitness Prediction.” Synthetic Biology. Available at: 10.1101/2023.12.07.570727.

Pei, H. et al. (2023) “Identification of Thermophilic Proteins Based on Sequence-Based Bidirectional Representations from Transformer-Embedding Features,” Applied Sciences, 13(5), p. 2858. Available at: 10.3390/app13052858.

Pourmirzaei, Mahdi et al. (2024) “Prot2Token: A multi-task framework for protein language processing using autoregressive language modeling.” ICML 2024 Workshop on Efficient and Accessible Foundation Models for Biological Discovery. Available at: https://openreview.net/forum?id=CxFKMxQA7f (Accessed: June 20, 2026).

Pudžiuvelytė, I., et al. (2024) “TemStaPro: protein thermostability prediction using sequence representations from protein language models,” Bioinformatics. Edited by L. Cowen, 40(4), p. btae157. Available at: 10.1093/bioinformatics/btae157.

Qing, R. et al. (2022) “Protein Design: From the Aspect of Water Solubility and Stability,” Chemical Reviews, 122(18), pp. 14085–14179. Available at: 10.1021/acs.chemrev.1c00757.

Ramsundar, B. et al. (2015) “Massively Multitask Networks for Drug Discovery.” arXiv. Available at: 10.48550/arXiv.1502.02072.

Rives, A. et al. (2021) “Biological structure and function emerge from scaling unsupervised learning to 250 million protein sequences,” Proceedings of the National Academy of Sciences, 118(15), p. e2016239118. Available at: 10.1073/pnas.2016239118.

Rosace, A. et al. (2023) “Automated optimisation of solubility and conformational stability of antibodies and proteins,” Nature Communications, 14(1), p. 1937. Available at: 10.1038/s41467-023-37668-6.

Schmirler, R., Heinzinger, M. and Rost, B. (2024) “Fine-tuning protein language models boosts predictions across diverse tasks,” Nature Communications, 15(1), p. 7407. Available at: 10.1038/s41467-024-51844-2.

Sledzieski, S. et al. (2024) “Democratizing protein language models with parameter-efficient fine-tuning,” Proceedings of the National Academy of Sciences, 121(26), p. e2405840121. Available at: 10.1073/pnas.2405840121.

Su, J. et al. (2023) “SaProt: Protein Language Modeling with Structure-aware Vocabulary.” Bioinformatics. Available at: 10.1101/2023.10.01.560349.

Thumuluri, V., et al. (2022) “NetSolP: predicting protein solubility in *Escherichia coli* using language models,” Bioinformatics. Edited by A. Valencia, 38(4), pp. 941–946. Available at: 10.1093/bioinformatics/btab801.

Unsal, S. et al. (2022) “Learning functional properties of proteins with language models,” Nature Machine Intelligence, 4(3), pp. 227–245. Available at: 10.1038/s42256-022-00457-9.

Wang, D. et al. (2025) “AI4Protein: transforming the future of protein design,” Science China Life Sciences, 68(10), pp. 2880–2890. Available at: 10.1007/s11427-024-2906-3.

Wang, H. et al. (2025) “DeepPFP: a multi-task-aware architecture for protein function prediction,” Briefings in Bioinformatics, 26(1), p. bbae579. Available at: 10.1093/bib/bbae579.

Zhou, Z. et al. (2024) “Enhancing efficiency of protein language models with minimal wet-lab data through few-shot learning,” Nature Communications, 15(1), p. 5566. Available at: 10.1038/s41467-024-49798-6.

Wolf, T. et al. (2020) “Transformers: State-of-the-art natural language processing,” in proceedings of the 2020 Conference on Empirical Methods in Natural Language Processing: System Demonstrations. Online: Association for Computational Linguistics, pp. 38-45. Available at: https://www.aclweb.org/anthology/2020.emnlp-demos.6

